# A Grid-Search Framework for Dataset-Specific Calibration of Actigraphy Sleep Detection Algorithms

**DOI:** 10.64898/2026.04.07.706161

**Authors:** Ali Rahjouei

## Abstract

Actigraphy is widely used for long-term sleep monitoring, but established sleep–wake scoring algorithms often require parameter tuning, which is commonly performed manually and can reduce reproducibility. In this study, a grid-search-based calibration framework is presented for established actigraphy algorithms and evaluate whether it can serve as a practical alternative to manual tuning. The method was evaluated using two datasets: a multi-subject polysomnography-validated actigraphy dataset and a self-collected dual-device dataset. In the polysomnography-validated dataset, grid-search optimization produced performance patterns similar to manual parameter selection, while slightly improving detection of sleep onset and sleep offset and yielding modest gains in wake-sensitive metrics. In the dual-device dataset, consensus and majority voting were useful for reducing the influence of brief wake episodes occurring within the main sleep period, including micro-awakenings that can fragment sleep predictions across individual algorithms. Overall, these findings show that grid-search can replace manual parameter tuning with a more explicit and reproducible procedure while providing small improvements in sleep timing estimation and benefiting ensemble-based handling of within-sleep wakefulness.

## 1. Introduction

Actigraphy, the continuous measurement of physical activity using wearable accelerometers, has become a cornerstone tool in sleep research and clinical practice. Unlike polysomnography (PSG), which requires specialized equipment and laboratory settings, actigraphy enables long-term, non-invasive monitoring of sleep patterns in naturalistic environments. Consequently, actigraphy is widely used in large-scale epidemiological studies, psychiatric research, and longitudinal monitoring of circadian rhythms (Fekedulegn et al., 2020; Smith et al., 2018).

A fundamental step in actigraphy analysis is the classification of sleep and wake states from activity signals. Over the past decades, numerous rule-based algorithms have been developed for this purpose, including the Cole–Kripke, Sadeh, Oakley, MASDA, and Crespo methods (Cole et al., 1992; Crespo et al., 2012; Loock et al., 2021; NR Oakley, 1997; Sadeh et al., 1994). These algorithms differ in their mathematical formulations, temporal smoothing strategies, and assumptions regarding the relationship between movement and sleep–wake states. Each algorithm also includes one or more tunable parameters, such as activity thresholds or smoothing window lengths, which strongly influence classification outcomes (Blackwell et al., 2011; Quante et al., 2018).

A major practical challenge in actigraphy research is that optimal parameter settings vary substantially across devices, populations, and recording contexts (Cellini et al., 2013; Meltzer et al., 2012). Differences in sensor sensitivity, device placement, individual activity patterns, and recording environments mean that parameter values validated in one study often perform poorly when applied to new datasets (Hjorth et al., 2012). As a result, researchers frequently rely on manufacturer defaults or informal manual tuning, leading to inconsistent results and reduced reproducibility across studies (De Souza et al., 2003; Quante et al., 2018).

Existing approaches typically treat algorithm selection and parameter tuning as independent decisions, optimizing each algorithm separately. However, this practice neglects an important observation: classical actigraphy algorithms often exhibit complementary strengths and weaknesses. For example, temporally smoothed algorithms may accurately capture consolidated sleep periods but fail to detect brief awakenings (Sadeh, 2011; Sadeh et al., 1994), whereas threshold-based algorithms may detect short wake bouts but produce fragmented sleep estimates. Consequently, selecting a single algorithm or tuning it in isolation may introduce systematic bias.

A systematic calibration framework is proposed in this work to address this problem, harmonizing multiple classical sleep detection algorithms for a given dataset and individual. Rather than developing a new predictive model, the goal is to automate the process of dataset-specific parameter tuning while reducing researcher degrees of freedom (Simmons et al., 2011).

The central idea explored in this work is that agreement between diverse sleep detection algorithms can be used as a practical criterion for selecting stable parameter configurations in the absence of labeled training data. Because classical actigraphy algorithms exhibit high parameter sensitivity—where small deviations can shift detection toward implausible extremes such as nearly continuous sleep or wake—solutions that produce consistent predictions across multiple algorithms are likely to reflect stable behavioral patterns. Rather than assuming that algorithm agreement represents ground truth, cross-algorithm convergence is used as an optimization target within physiological plausibility constraints. This allows automated calibration of algorithm parameters while reducing researcher degrees of freedom.

Since individual sleep detection algorithms exhibit high parameter sensitivity, where minor deviations can drastically shift detection towards non-physiological extremes (0% or 100% sleep), it is suggested as highly probable that the true, stable behavioral pattern is captured when five diverse algorithms reach a simultaneous alignment in their parameter configurations. Therefore, by optimizing for maximum inter-algorithm agreement within physiological constraints, the identification of accurate detection parameters is achieved without reliance on labeled training data. To operationalize this idea, a grid-search-based calibration procedure was implemented to explore parameter combinations across multiple algorithms. The optimization balances several criteria: maximizing inter-algorithm agreement, enforcing physiological plausibility constraints, minimizing temporal instability, and preventing misclassification of non-wear periods as sleep. The calibrated algorithm outputs are then integrated through ensemble decision rules that quantify uncertainty and allow abstention in low-confidence situations.

The proposed framework is evaluated using two datasets: a dual-device self-recorded dataset and a polysomnography-validated actigraphy dataset (Walch et al., 2019). These findings show that automated calibration of research-grade actigraphy can tune algorithm parameters to produce sleep bout estimates that align with those derived from a commercial smartwatch optimized for its own sensor data (“Estimating Sleep Stages from Apple Watch,” 2025). Furthermore, applying ensemble methods, specifically majority voting and consensus region identification, helps reduce fragmentation caused by brief wake episodes within the main sleep period. However, comparison with PSG highlights an intrinsic limitation of activity-based sleep detection: algorithmic convergence reflects behavioral quiescence rather than electrophysiologically defined sleep.

## Contributions

This work makes four primary contributions:

**Calibration Framework:** a systematic grid-search approach for dataset-specific calibration of classical actigraphy sleep detection algorithms.

**Automated Parameter Selection:** demonstration that automated calibration can replace manual parameter tuning while producing comparable performance and slightly improved sleep timing estimates.

**Ensemble-Based Sleep Detection:** evaluation of consensus and majority voting approaches for reducing fragmentation caused by brief wake episodes within the main sleep period.

**Insight into Actigraphy Limitations:** comparison with PSG highlights that actigraphy algorithms primarily capture behavioral quiescence rather than electrophysiologically defined sleep.

## 2. Methods

### 2.1 Study Design and Datasets

The proposed framework is designed as a **dataset-specific calibration pipeline** for harmonizing classical actigraphy sleep detection algorithms. Rather than training a predictive model, the method optimizes algorithm parameters independently for each recording to account for device characteristics, individual activity patterns, and recording conditions.

The framework was evaluated using two datasets with distinct purposes.

#### 2.1.1 Dataset 1: Polysomnography-Validated Actigraphy Dataset

A multi-subject dataset containing concurrent wrist-worn actigraphy and polysomnography (PSG) recordings was used to evaluate calibrated algorithm outputs against physiological sleep staging. Activity signals were recorded at 1-second resolution and later aggregated to 1-minute epochs.

PSG sleep stages were binarized into wake (stage 0) and sleep (stages N1, N2, N3, REM) to provide a reference standard for epoch-level comparison (Walch et al., 2019).

#### 2.1.2 Dataset 2: Dual-Device Self-Recording Dataset

A self-recorded dataset was collected to develop and illustrate the calibration pipeline. A single participant simultaneously wore a research-grade actigraph (ActiLumus) and an Apple Watch on opposite wrists for ten consecutive days.

Apple Watch sleep-stage outputs were binarized (sleep stage > 0 = sleep) and used as an external reference for algorithm comparison. This dataset was used for qualitative evaluation of multi-day algorithm behavior, rather than for generalizable validation.

### 2.2 Data Preprocessing

All datasets underwent standardized preprocessing to ensure consistency across recordings.

#### 2.2.1 Temporal Alignment and Epoch Standardization

Activity data were resampled to uniform 60-second epochs. For high-resolution signals, missing timestamps were first interpolated to create continuous timelines before aggregation.

#### 2.2.2 Complete-Day Filtering

Only complete recording days containing exactly 1,440 epochs were retained to avoid edge effects in circadian analyses.

### 2.3 Non-Wear Detection

Off-wrist periods were identified using an adapted Choi algorithm based on extended zero-activity runs with artifact tolerance (Choi et al., 2012). Continuous low-activity segments exceeding a minimum duration threshold were classified as non-wear.

### 2.4 Activity Imputation

A two-stage imputation procedure replaced activity values during non-wear periods using time-of-day baseline activity patterns (dataset 1 and 2). For continuous activity metrics, imputation used percentile-bounded random sampling to preserve realistic signal distributions (dataset 2).

### 2.5 manual adjustment

For comparison with the automated calibration framework, each algorithm was also evaluated using manually selected parameter settings. In this configuration, parameter values were adjusted iteratively while visually inspecting the resulting sleep–wake classifications alongside the raw activity signal.

Parameter combinations were modified until the detected sleep periods aligned with extended intervals of reduced activity that visually resembled consolidated sleep bouts. This procedure reflects common practice in exploratory actigraphy analyses, where parameter settings are often adjusted to match observable activity patterns in the absence of a physiological ground truth.

Because actigraphy is frequently used in long-term behavioral monitoring without concurrent polysomnography, this manual tuning approach provides a practical baseline representing typical real-world usage conditions. The manually selected configurations were therefore used as a reference for evaluating the potential benefits of the proposed automated calibration framework.

### 2.6 Performance Evaluation

Performance metrics included accuracy, balanced accuracy, precision, recall, specificity, F1-score, Cohen’s kappa (Cohen, 1960), and Matthews correlation coefficient (MCC). MCC and balanced accuracy were prioritized to account for the inherent class imbalance between sleep and wake states in actigraphy data (Chicco & Jurman, 2020). Importantly, these metrics were used to characterize calibrated outputs rather than to train predictive models, ensuring an independent assessment of the harmonization framework’s efficacy.

### 2.7 Implementation and Reproducibility

The sleep-wake scoring algorithms used in this study were implemented based on their original published descriptions and validated against the open-source pyActigraphy software package (Hammad et al., 2021). The present work does not introduce novel implementations of these algorithms; rather, the methodological contribution lies in the proposed multi-objective calibration and harmonization framework.

### 2.8 Sleep Onset and Offset Detection and Evaluations

Sleep onset and offset times were derived from the binary sleep–wake sequences produced by each algorithm and by the reference labels. For each subject, records were ordered by timestamp and short wake interruptions of ≤10 consecutive minutes occurring within sleep were bridged to reduce the influence of brief awakenings. After this gap-filling step, the main sleep episode was defined as the longest contiguous run of sleep epochs in the recording. Sleep onset was taken as the first epoch of this main sleep bout, and sleep offset as the last epoch of the same bout.

Algorithm-derived onset and offset times were compared with the corresponding reference times obtained from PSG (Dataset 1) or Apple Watch-derived binary labels (Dataset 2). Timing errors were calculated in minutes as algorithm estimate minus reference estimate, such that positive values indicate a later estimated event and negative values indicate an earlier estimated event.

Agreement was evaluated using error distributions, mean error (ME), mean absolute error (MAE), root mean square error (RMSE), scatter plots against the identity line, and Bland–Altman analysis.

## 3 Grid-Search workflow and rationale

The proposed calibration framework was applied to identify parameter configurations for multiple classical actigraphy sleep–wake algorithms that produce physiologically plausible and mutually consistent sleep classifications. Rather than optimizing algorithm parameters against a labeled reference dataset, the framework uses a consensus-based optimization strategy, selecting parameter configurations that maximize agreement between algorithms while enforcing physiological plausibility constraints.

The calibration procedure consisted of three main stages: candidate parameter filtering, consensus-based optimization, and selection of the final parameter configuration (Fig. 1). These stages progressively reduced the parameter search space and identified a parameter combination that yielded consistent sleep–wake classifications across algorithms.

**Figure 1.**
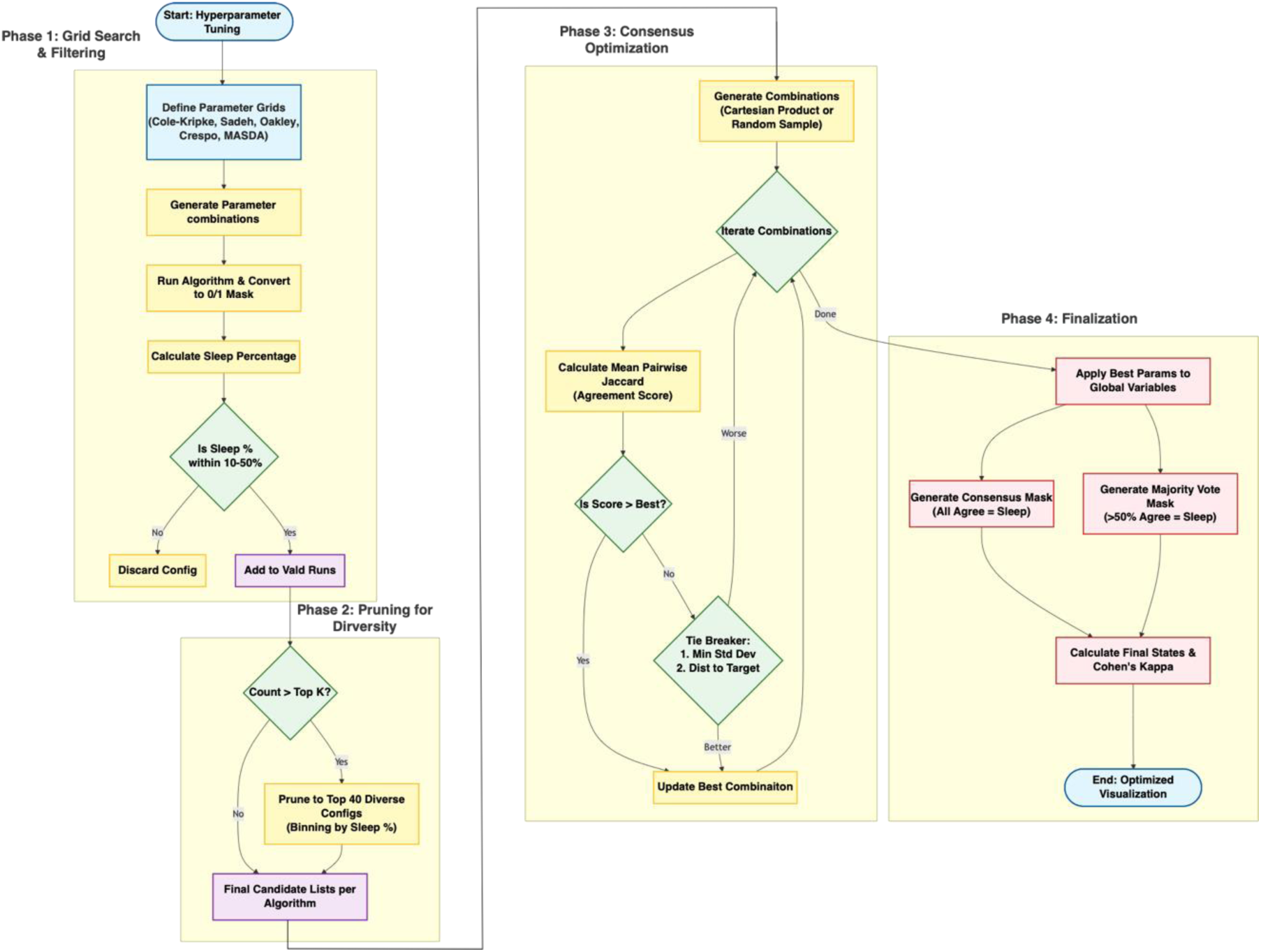
Hyperparameter tuning and consensus optimization workflow for actigraphy-based sleep detection algorithms. The process is divided into four distinct phases: **(Phase 1)** Grid Search & Filtering, where parameter grids are defined for five algorithms (Cole-Kripke, Sadeh, Oakley, Crespo, MASDA) and results are filtered to retain only biologically plausible sleep percentages (10–50%); **(Phase 2)** Pruning for Diversity, which reduces the computational load by selecting the top 𝐾 (e.g., 40) configurations distributed across sleep percentage bins; **(Phase 3)** Consensus Optimization, which iterates through parameter combinations to maximize inter-algorithm agreement (Mean Pairwise Jaccard Index), using sleep duration standard deviation as a tie-breaker; and **(Phase 4)** Finalization, where the optimal parameters are applied to generate strict consensus (100% agreement) and majority vote (>50% agreement) sleep masks, validated by Cohen’s Kappa statistics.

### 3.1 Candidate Parameter Filtering

The calibration process began with a broad grid search over candidate parameter values for each algorithm. The evaluated algorithms included Cole–Kripke, Sadeh, Oakley, Crespo, and MASDA (Roenneberg), each of which contains parameters controlling classification thresholds, temporal smoothing, or activity weighting.

For each algorithm, a predefined parameter grid was constructed spanning a range of plausible values based on algorithm design and prior usage. Each parameter combination was applied to the activity signal to produce a binary sleep–wake mask, representing epochs classified as sleep or wake. Because some parameter configurations can produce unrealistic sleep estimates, a physiological plausibility filter was applied to remove implausible results. Specifically, configurations were retained only if the proportion of epochs classified as sleep fell within a predefined range of 10–50 % of the recording duration. This filtering step eliminated parameter settings that yielded extreme outcomes such as near-continuous sleep or almost no detected sleep. Even after this filtering stage, the number of valid parameter configurations could remain large. To ensure computational feasibility in the subsequent optimization stage, candidate configurations were further reduced by retaining a diverse subset of valid runs for each algorithm. Diversity was assessed based on predicted sleep percentages and mask variability, ensuring that the remaining candidates covered a broad range of plausible behaviors while limiting the total number of configurations.

This stage therefore produced a manageable set of candidate parameterizations for each algorithm while ensuring that all retained configurations generated physiologically plausible sleep estimates.

### 3.2 Consensus-Based Optimization

After identifying valid parameter candidates for each algorithm, the next stage aimed to determine which combination of parameter settings across algorithms produced the highest agreement in sleep–wake classification. Candidate configurations from all algorithms were combined to form multi-algorithm parameter sets. For each combination, agreement between the resulting sleep masks was quantified using the mean pairwise Jaccard similarity, which measures the overlap between binary sleep–wake sequences. High Jaccard similarity indicates that multiple algorithms classify the same epochs as sleep or wake, suggesting greater consistency in sleep detection.

The primary optimization objective was to maximize the mean pairwise Jaccard similarity across all algorithms. This approach treats agreement between established algorithms as a proxy for reliable classification when an external ground truth is unavailable.

When multiple parameter combinations produced identical agreement scores, two secondary selection criteria were applied to break ties:

1. **Consistency of predicted sleep duration.** Preference was given to parameter combinations that minimized the standard deviation of predicted sleep percentages across algorithms, ensuring that different algorithms estimated similar total sleep durations.
2. **Physiological plausibility of sleep duration.** Solutions were favored when the mean predicted sleep percentage across algorithms was closer to a target value representing typical nightly sleep proportions.

In some cases, the total number of possible parameter combinations across algorithms exceeded practical computational limits. When this occurred, combinations were evaluated using random sampling of the parameter space, allowing the optimization to explore a large number of candidate configurations without exhaustive enumeration.

Through this consensus-based optimization process, the framework identified parameter combinations that produced the most mutually consistent sleep–wake classifications across algorithms while maintaining physiologically plausible sleep estimates.

### 3.3 Selected Parameter Configurations

The parameter combination achieving the highest consensus score was selected as the **final calibrated configuration** for each algorithm. These optimized parameters were then applied to generate the final binary sleep–wake masks for all algorithms.

Using these calibrated parameters, two ensemble sleep-wake masks were additionally constructed:

- **Strict consensus mask**, where an epoch was labeled as sleep only if all algorithms agreed.
- **Majority voting mask**, where an epoch was labeled as sleep when more than half of the algorithms classified the epoch as sleep.

These ensemble masks provide complementary perspectives on sleep detection. The strict consensus mask emphasizes high confidence classifications with maximal agreement, whereas the majority voting mask offers a more permissive estimate of sleep periods.

The calibrated algorithm outputs and ensemble masks served as the basis for the subsequent performance evaluations. In the following sections, these calibrated classifications are compared with reference sleep labels derived from polysomnography recordings (Section 4) and manual parameter configurations (Section 5), followed by analyses of sleep timing agreement (Section 6) and multi-day algorithm behavior (Section 7).

## 4. Results

### 4.1 Algorithm performance with manual parameters

Across the 23 subjects, all algorithms demonstrated high sensitivity for sleep detection, with recall values approaching unity, indicating that the majority of PSG-defined sleep epochs were correctly classified as sleep (**Fig. 2A–B**). However, specificity for wake detection remained lower, reflecting frequent misclassification of wake epochs as sleep.

**Figure 2.**
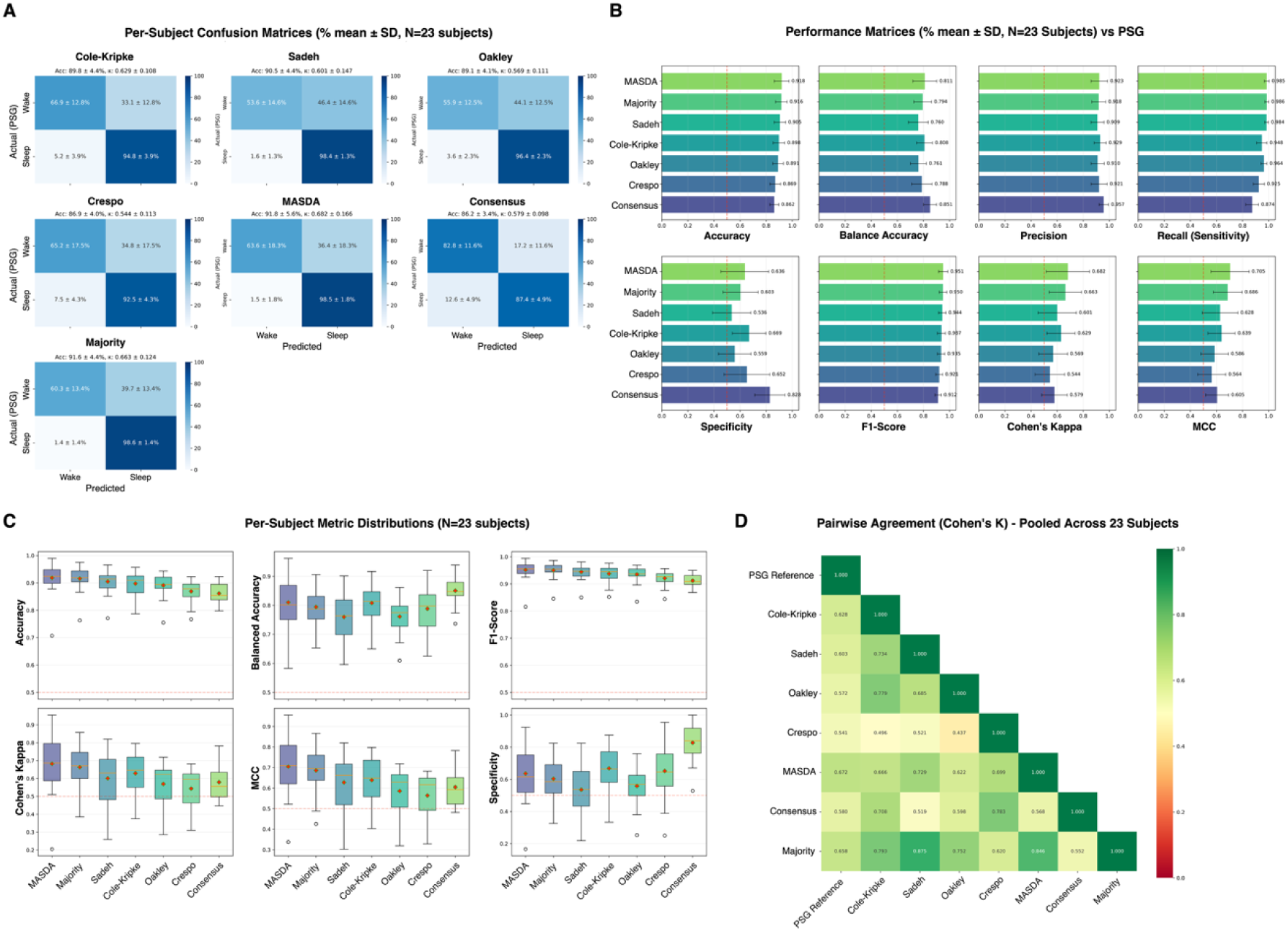
Performance of sleep–wake algorithms using manually selected parameter settings across 23 subjects. **(A)** Mean per-subject confusion matrices (±SD) relative to the PSG reference. Values represent the percentage of epochs classified as wake or sleep averaged across subjects. Across algorithms, sleep epochs were identified with high sensitivity, whereas wake epochs were more frequently misclassified as sleep. **(B)** Mean performance metrics (±SD) across subjects, including Accuracy, Balanced Accuracy, Precision, Recall (Sensitivity), Specificity, F1-score, Cohen’s κ, and Matthews Correlation Coefficient (MCC). The red dashed lines indicate reference thresholds for comparison across metrics. **(C)** Boxplots showing the distribution of performance metrics across subjects (N = 23). While accuracy and F1-score remain consistently high across algorithms, greater variability is observed in specificity and agreement-based metrics such as Cohen’s κ and MCC. **(D)** Pairwise agreement heatmap showing Cohen’s κ values between the PSG reference and each algorithm, as well as agreement among the algorithms themselves. Higher agreement values indicate greater similarity in sleep–wake classification patterns.

Overall accuracy and F1-scores were relatively high across algorithms, largely driven by the high sensitivity to the dominant sleep class (**Fig. 2B**). In contrast, metrics that account for class imbalance and chance agreement, including balanced accuracy, Cohen’s κ, and Matthews Correlation Coefficient (MCC), were comparatively lower. This indicates that the algorithms were less effective in identifying wake periods during the sleep interval. Analysis of per-subject metric distributions revealed consistent patterns across the cohort (**Fig. 2C**). Sensitivity showed limited variability across subjects, whereas specificity and agreement-based metrics exhibited greater dispersion, suggesting that wake detection performance varied more substantially between individuals. Finally, pairwise agreement analysis demonstrated moderate agreement between algorithms and the PSG reference, as well as among the algorithms themselves (**Fig. 2D**). These results indicate that while the algorithms produce broadly similar classification patterns, their agreement with the physiological sleep reference remains constrained by difficulties in detecting wake epochs.

### 4.2 Algorithm performance with grid-search optimization

For each algorithm, parameters were optimized using a custom grid search framework prior to evaluation as described previously. Across the 23 subjects, all algorithms demonstrated high sensitivity for sleep detection, with mean recall values approaching unity (≈0.97–1.00). This indicates that the majority of PSG-defined sleep epochs were correctly classified as sleep by the algorithms.

However, specificity for wake detection was substantially lower, reflecting frequent misclassification of wake epochs as sleep in the pooled confusion matrices (**Fig. 3A**).

**Figure 3.**
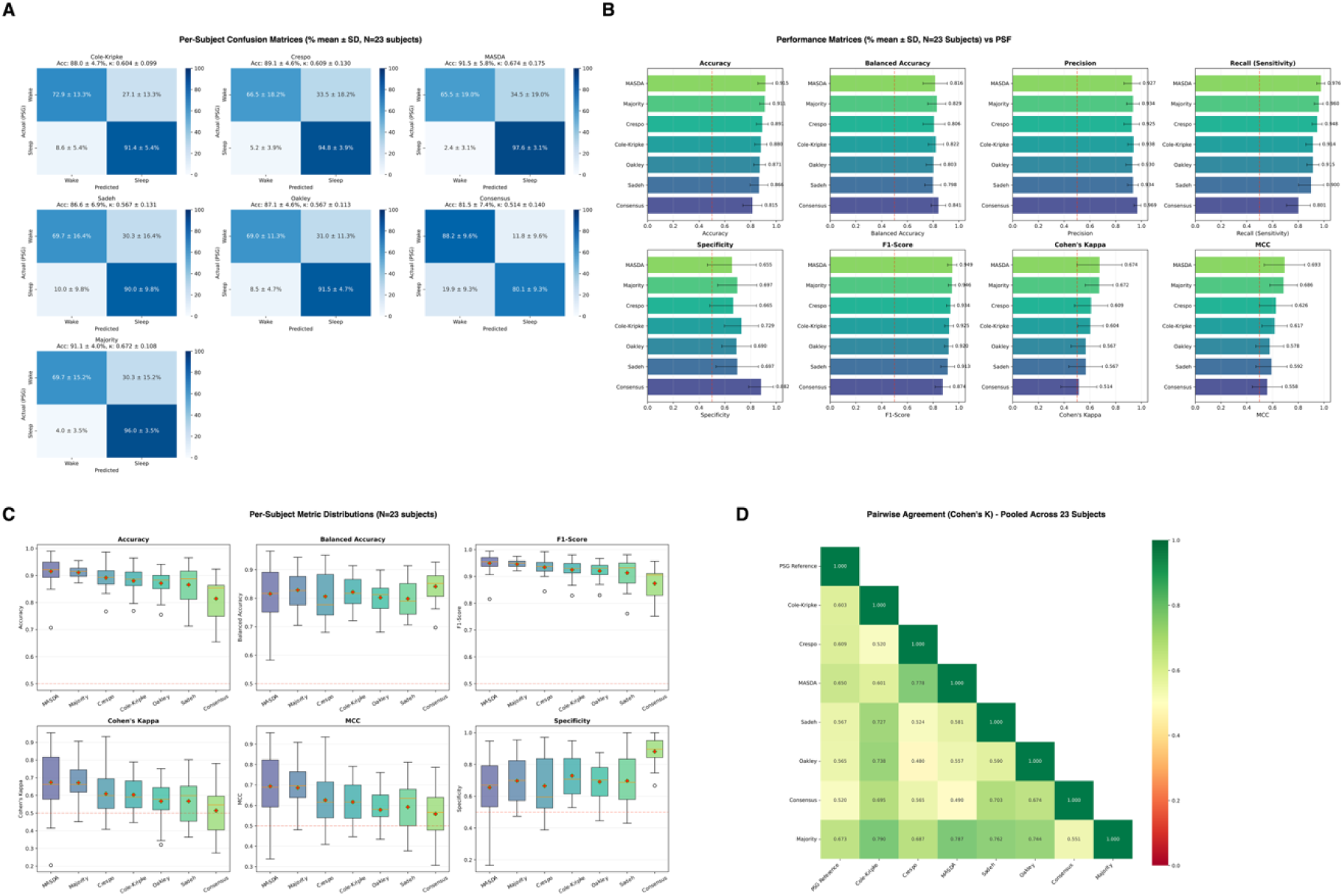
Performance of actigraphy-based sleep–wake algorithms compared with polysomnography (PSG) across 23 subjects. **(A)** Pooled confusion matrices showing epoch-level classification relative to PSG for each algorithm. **(B)** Mean performance metrics across subjects (±SD), including Accuracy, Balanced Accuracy, Precision, Recall (Sensitivity), Specificity, Cohen’s κ, F1-score, and Matthews Correlation Coefficient (MCC). **(C)** Boxplots showing per-subject distributions of performance metrics across the cohort (N = 23). Red dashed lines indicate commonly used reference thresholds for agreement (e.g., κ = 0.6). **(D)** Heatmap of pairwise Cohen’s κ values showing agreement between algorithms and PSG.

Despite relatively high overall accuracy and F1-scores, metrics that account for class imbalance and chance agreement, including balanced accuracy, Cohen’s κ, and Matthews Correlation Coefficient (MCC), were noticeably lower (**Fig. 3B**). This pattern reflects the imbalance between sleep and wake epochs in overnight recordings, where strong performance on the dominant sleep class can inflate conventional performance metrics.

Analysis of per-subject metric distributions showed that this pattern was consistent across the cohort (**Fig. 3C**). Sensitivity remained tightly clustered near its upper bound, whereas specificity, balanced accuracy, κ, and MCC exhibited greater variability between subjects.

Finally, pairwise agreement analysis using Cohen’s κ demonstrated moderate agreement between the different algorithms and PSG, as well as among the algorithms themselves (**Fig. 3D**), indicating that while the algorithms generate broadly similar classification patterns, their agreement with the physiological reference remains constrained by difficulties in identifying wake epochs during sleep periods.

### 4.3 Comparison between tuning approaches

To evaluate the impact of parameter selection on algorithm performance, results obtained using manually selected parameter configurations (**Fig. 2**) were compared with those obtained using grid-search optimized parameters (**Fig. 3**). Overall performance patterns were broadly similar between the two approaches. Across all algorithms, sleep detection remained highly sensitive, with recall values consistently approaching unity under both parameter configurations. This indicates that both tuning strategies preserved strong detection of PSG-defined sleep epochs. However, differences emerged in metrics reflecting wake detection and overall classification balance. Grid-search optimization generally resulted in modest improvements in balanced accuracy, specificity, and agreement-based metrics, including Cohen’s κ and Matthews Correlation Coefficient (MCC). These improvements suggest that automated parameter optimization provided better calibration of algorithm thresholds, leading to improved discrimination between sleep and wake epochs. In contrast, manually selected parameters tended to maintain high overall accuracy and F1-scores but showed comparatively lower performance in metrics sensitive to class imbalance. Per-subject metric distributions further illustrated these differences. While both approaches demonstrated limited variability in sleep sensitivity across subjects, the grid-search optimized configuration showed slightly reduced variability in specificity and agreement-based metrics, indicating more consistent performance across the cohort. Despite these improvements, the overall classification patterns remained similar between the two parameterization strategies. Pairwise agreement analysis revealed that algorithms continued to exhibit moderate agreement with the PSG reference and comparable agreement with one another, regardless of the parameter tuning method.

Taken together, these findings indicate that automated grid-search optimization yields performance comparable to manually selected parameter configurations while providing a more systematic and reproducible approach to parameter tuning. Thus, grid-search optimization can serve as a practical replacement for manual parameter selection without altering the fundamental performance characteristics of actigraphy-based sleep–wake classification.

### 4.4 Sleep onset and offset agreement with PSG

Agreement between algorithm-derived sleep timing estimates and PSG was evaluated by comparing sleep onset and sleep offset times obtained using both manual parameter settings and grid-search optimized configurations (**Fig. 4**). Across algorithms, sleep onset and offset estimates generally followed the PSG reference, as illustrated by the clustering of points around the identity line in the scatter plots (**Fig. 4B**). However, systematic deviations were observed across subjects and algorithms. The distribution of timing errors indicated that sleep onset estimates tended to be closer to the PSG reference than sleep offset estimates, although variability was present across algorithms (**Fig. 4A**). Comparing parameterization strategies, the grid-search optimized configurations generally reduced timing errors and produced estimates that more closely aligned with PSG, particularly for sleep onset. This improvement was reflected by tighter clustering of points around the identity line in the scatter plots and narrower error distributions across subjects. In contrast, manually selected parameters exhibited larger deviations from PSG for some algorithms. Bland–Altman analyses further characterized the agreement patterns (**Fig. 4C**). Across algorithms, the mean bias between algorithm-derived estimates and PSG varied depending on the algorithm and parameter configuration. Grid-search optimization generally resulted in reduced bias and narrower limits of agreement, indicating improved consistency of sleep timing estimates across subjects. Nevertheless, the limits of agreement remained relatively wide for several algorithms, highlighting persistent variability in estimating sleep onset and offset from actigraphy data. Together, these results indicate that while actigraphy-based algorithms can approximate PSG-derived sleep timing, parameter optimization through grid search improves agreement and reduces variability, particularly for sleep onset estimation.

**Figure 4.**
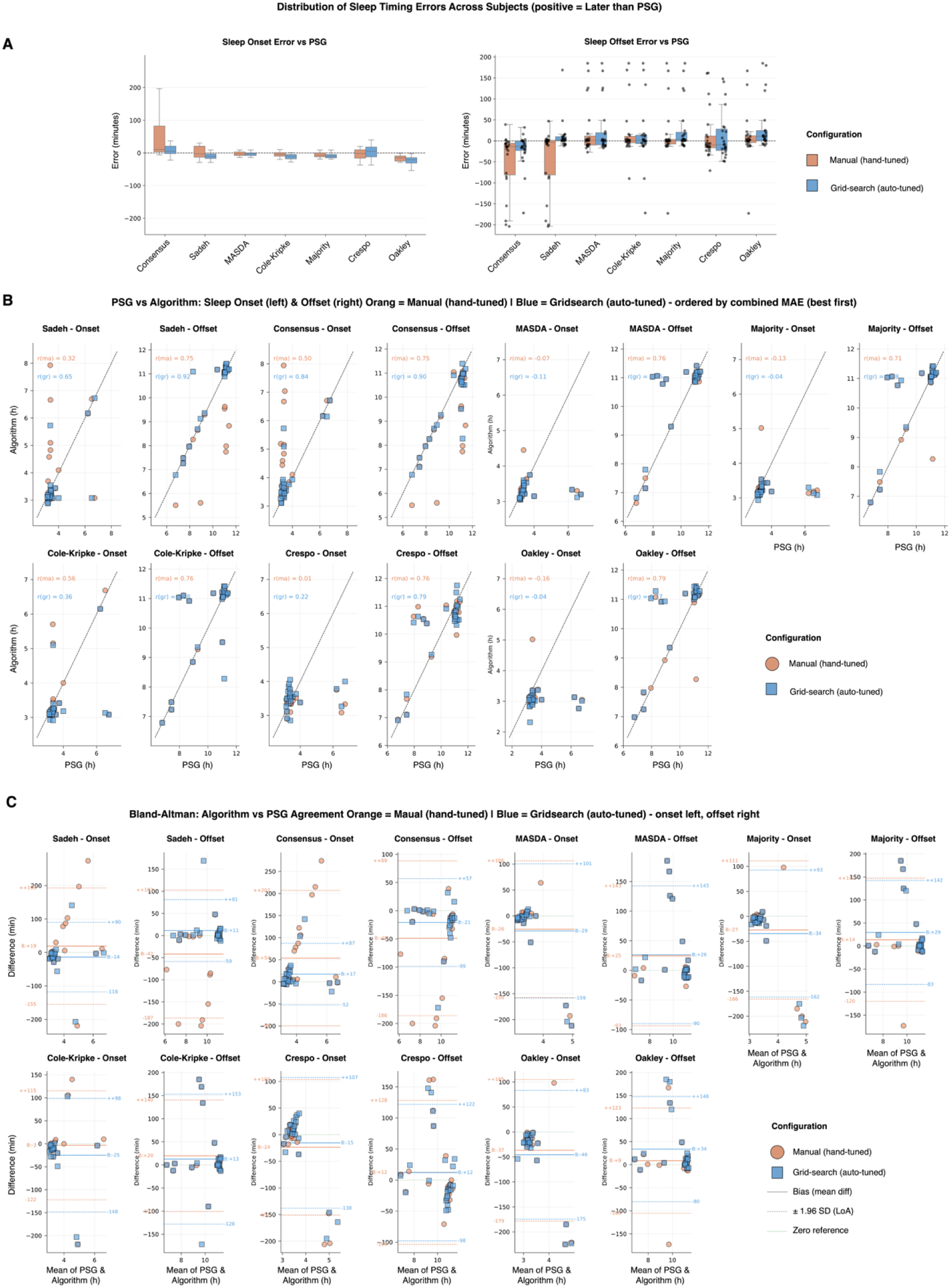
Comparison of sleep timing estimates between algorithms and PSG using manual and grid-search parameter configurations. **(A)** Distribution of sleep onset and sleep offset timing errors across subjects relative to PSG (N = 23). Positive values indicate estimates occurring later than the PSG reference, while negative values indicate earlier estimates. Results are shown for both manual parameter settings (orange) and grid-search optimized parameters (blue). **(B)** Scatter plots comparing algorithm-derived sleep onset (left panels) and sleep offset (right panels) times against PSG measurements for each algorithm. Points represent individual subjects. The diagonal dashed line represents perfect agreement with PSG. Manual parameter results are shown in orange, and grid-search optimized results in blue. Algorithms are ordered according to the combined mean absolute error (MAE). **(C)** Bland–Altman plots illustrating agreement between algorithm estimates and PSG for sleep onset (left panels) and sleep offset (right panels). Each point represents a subject. The central line indicates the mean difference (bias) between algorithm and PSG, and the dashed lines represent the limits of agreement (±1.96 SD). Results for manual parameter settings are shown in orange, while grid-search optimized parameters are shown in blue.

### 4.5 Multi-Day Sleep Detection and Fragmentation Analysis

To investigate algorithm performance under longitudinal conditions, multi-day actigraphy data from a single subject were analyzed (Fig. 5). Due to limitations in continuous PSG recordings, the subject wore both a research-grade actigraph and an Apple Watch Series 10 simultaneously, allowing sleep staging data from the smartwatch to serve as an external reference. Apple Watch sleep stage outputs were converted to binary sleep–wake labels (sleep = 1, wake = 0) for direct comparison with algorithm predictions.

**Figure 5.**
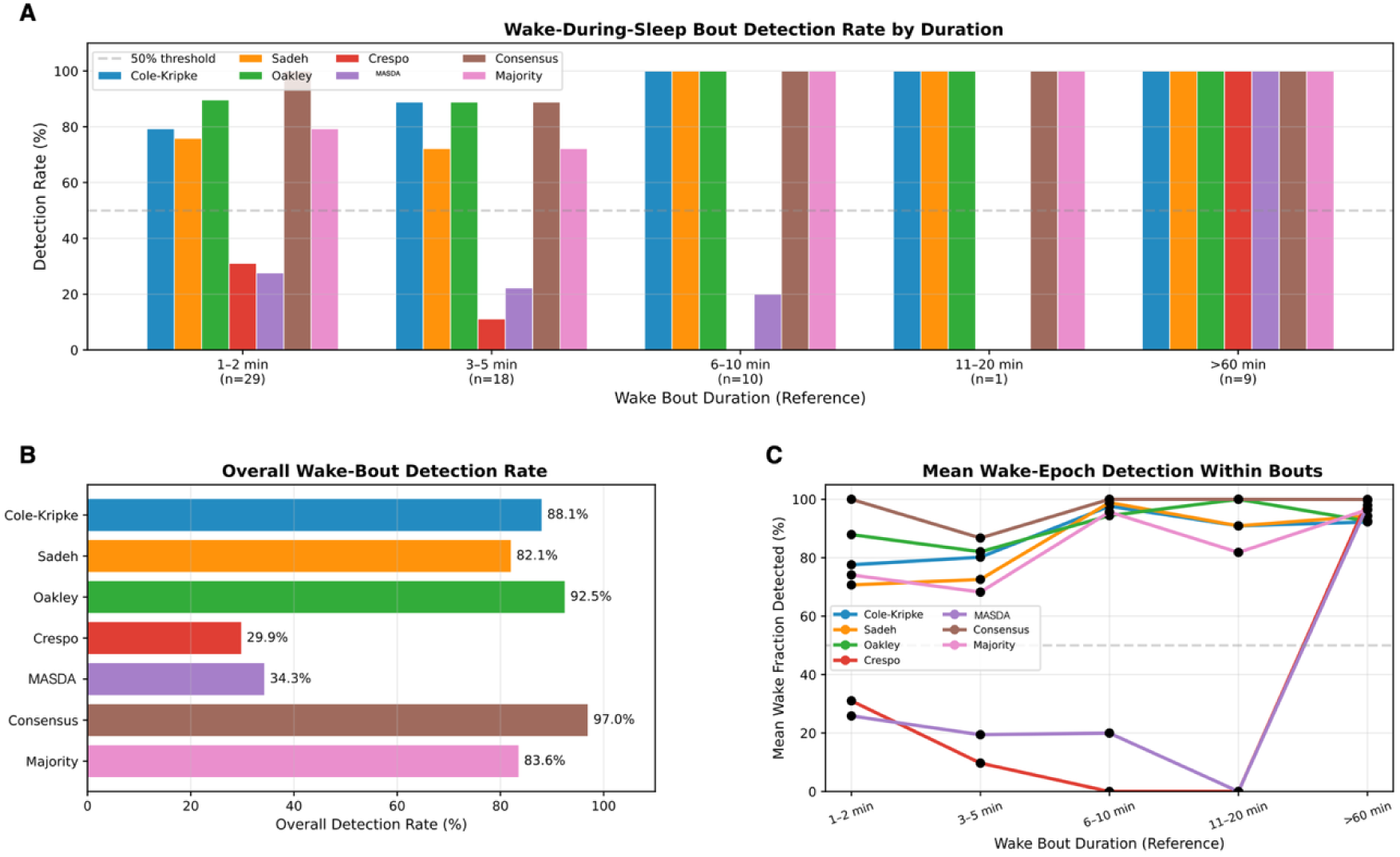
Detection of wake bouts occurring within sleep across actigraphy algorithms. **(A)** Wake-bout detection rate by reference bout duration. Wake bouts were defined from the reference sleep–wake signal as contiguous runs of wake occurring within sleep (i.e., flanked by sleep on both sides). A bout was considered detected when an algorithm classified ≥50% of epochs within the bout as wake. Detection rates are shown for duration bins (1–2 min, 3–5 min, 6–10 min, 11–20 min, >60 min). Sample sizes for each bin are indicated below the x-axis labels. Colors represent algorithms: Cole–Kripke, Sadeh, Oakley, Crespo, MASDA, Consensus, and Majority. The dashed horizontal line indicates the 50% detection threshold. **(B)** Overall wake-bout detection rate across all durations. Values represent the percentage of reference wake bouts correctly detected by each algorithm. **(C)** Mean fraction of wake epochs detected within each bout, averaged by duration bin. This metric reflects how much of each wake bout is identified as wake by the algorithm, independent of the binary detection threshold.

Across multiple consecutive days, the algorithms using grid-search optimized parameters were able to reproduce the overall sleep–wake structure observed in the reference signal (Supp. 3A–B). Major sleep periods were consistently identified, and transitions between wake and sleep generally aligned with the smartwatch-derived labels. Pairwise Cohen’s κ analysis showed that grid-search–optimized algorithms exhibited higher agreement both with each other and with the smartwatch-derived labels compared with PSG-based comparisons (Supp. 3C).

Wake bouts occurring within sleep were identified in the reference signal and grouped by duration to evaluate how well each algorithm detected brief awakenings (Fig. 5A). Short bouts were the most common, with 29 bouts lasting 1–2 min and 18 lasting 3–5 min. Detection performance varied substantially for these short awakenings. For 1–2 min bouts, traditional algorithms such as Cole–Kripke, Sadeh, and Oakley detected approximately 75–90% of events, whereas Crespo and MASDA detected substantially fewer (∼30% and ∼27%, respectively). The ensemble approaches performed strongly, with Consensus detecting all 1–2 min bouts and the Majority method detecting ∼80%. A similar pattern was observed for 3–5 min bouts, where detection remained high for Cole–Kripke, Oakley, and Consensus (∼88–90%), moderate for Sadeh (∼72%), and markedly lower for Crespo (∼11%) and MASDA (∼22%).

Detection rates increased with bout duration. For bouts lasting 6–10 min or longer, most algorithms—including Cole–Kripke, Sadeh, Oakley, Consensus, and Majority—detected nearly all events. Crespo also detected longer bouts reliably, while MASDA continued to detect only a small fraction of 6–10 min bouts. Across all durations combined (Fig. 5B), Consensus achieved the highest overall detection rate (97.0%), followed by Oakley (92.5%), Cole–Kripke (88.1%), Majority (83.6%), and Sadeh (82.1%), whereas Crespo (29.9%) and MASDA (34.3%) detected far fewer wake bouts. Consistent with these patterns, the mean fraction of wake epochs detected within bouts (Fig. 5C) was high for Cole–Kripke, Sadeh, Oakley, and Consensus across most duration bins (generally ∼80–100%), while Crespo and MASDA captured substantially smaller portions of wake epochs during short awakenings.

## 5. Discussion

The main contribution of this study is demonstrating that classical actigraphy sleep–wake algorithms can be calibrated simultaneously using an unsupervised, consensus-based grid-search framework without compromising performance relative to manual parameter adjustment. This addresses a practical challenge in wearable sleep research: many real-world studies, particularly long-term behavioral monitoring, lack physiological references such as PSG, yet investigators must still select plausible and reproducible algorithm parameters. Current sleep medicine guidance supports the use of actigraphy but emphasizes that interpretation depends strongly on the scoring approach and context of use (Smith et al., 2018).

A central finding is that the grid-search strategy identified parameter sets that produced high agreement across algorithms while maintaining physiologically plausible sleep estimates. Traditional workflows often rely on default settings, proprietary software parameters, or manual tuning guided by visual inspection. Although practical, these approaches are difficult to standardize and reproduce across studies or raters. Even structured manual scoring introduces subjectivity and requires explicit procedures to achieve reproducibility (Breneman et al., 2024).

The proposed framework replaces subjective adjustment with an explicit optimization rule. Instead of relying on visual judgment of whether detected intervals “look like sleep,” the algorithm searches a predefined parameter space, removes implausible solutions, and selects the configuration that maximizes inter-algorithm agreement. Although the method still depends on assumptions about plausible sleep proportions and algorithm behavior, it makes the calibration procedure transparent, auditable, and repeatable. Future work will extend this approach by incorporating short-term activity averaging (e.g., 30–60 minute sliding windows) to detect sustained low-activity intervals that may correspond to resting periods.

Importantly, consensus does not imply validity. Algorithms may agree while sharing systematic biases. In PSG comparisons, the calibrated algorithms showed the typical actigraphy pattern of high sleep sensitivity but lower wake specificity, with moderate balanced accuracy, kappa, and MCC. This reflects a known limitation of wrist actigraphy rather than a failure of the framework. Numerous studies show that actigraphy detects sleep well but tends to overestimate sleep and underestimate wake, particularly quiet wakefulness and wake after sleep onset(Quante et al., 2018).

Thus, consensus-based calibration improves reproducibility and provides a principled way to tune parameters without labels but cannot overcome the fundamental sensing limitations of wrist accelerometry. Because accelerometers measure movement rather than neurophysiological sleep, immobile wakefulness remains difficult to distinguish from sleep. This explains why high recall and overall accuracy were accompanied by lower specificity and moderate chance-corrected agreement metrics, consistent with previous actigraphy studies (Quante et al., 2018).

This perspective also clarifies the comparison with manual tuning. In this study, manual parameter selection involved visually adjusting outputs until detected sleep periods appeared to align with low-activity intervals. While this incorporates contextual human judgment and can adapt to unusual recordings, it is difficult to standardize and vulnerable to operator bias. The grid-search framework offers a rule-based alternative that can be applied consistently across recordings. The fact that optimized configurations generally improved classification balance and timing agreement relative to manual tuning suggests that systematic parameter search can outperform ad hoc adjustment under the conditions examined here. Human scorers tend to prefer outputs that appear physiologically plausible and internally coherent; the framework operationalizes similar principles through explicit constraints and agreement criteria. This is particularly valuable in large-scale or technical studies where reproducibility is essential. The importance of subject-specific calibration is also supported by prior work showing that personalized models can outperform generalized ones in actigraphy-based sleep estimation (Khademi et al., 2019).

The ensemble results provide a second contribution. Combining calibrated algorithms through strict consensus and majority voting produced interpretable outputs. Ensemble approaches are attractive because classical actigraphy algorithms differ in their assumptions about thresholds, smoothing, and sleep continuity. Majority voting reduces the influence of individual algorithm biases, whereas strict consensus emphasizes high-confidence intervals where all algorithms agree. Ensemble strategies have repeatedly improved robustness and generalization in automated sleep scoring tasks.

These strategies involve different trade-offs. Strict consensus is conservative and may reduce false-positive sleep detection but can fragment sleep and increase false wake calls. Majority voting is more permissive and better preserves the continuity of the main sleep episode but may inherit actigraphy’s tendency to classify quiet wake as sleep. Accordingly, ensemble choice should depend on the application: majority voting may be preferable for estimating overall sleep windows, whereas strict consensus may better identify high-confidence sleep intervals.

Sleep onset and offset analyses further highlight where consensus-based optimization is beneficial. Grid-searched configurations generally aligned more closely with reference timing and reduced error relative to manual tuning, particularly for sleep onset. Because onset and offset represent transition points, parameter settings that improve agreement around transitions naturally improve timing estimates. Nevertheless, Bland–Altman analyses showed wide limits of agreement for some algorithms, reflecting persistent uncertainty in actigraphy-based timing estimation. This is consistent with literature indicating that actigraphy is more reliable for estimating overall sleep schedules and sleep opportunity than for precisely resolving sleep latency, wake after sleep onset, or fragmentation relative to PSG (Conley et al., 2019).

The multi-day single-subject analysis extends these findings to ecological monitoring, where actigraphy is often most valuable. Long-term PSG is rarely feasible, so wearable devices are frequently used for longitudinal sleep tracking. In this context, the ability to calibrate classical algorithms without PSG is particularly useful. The multi-day results suggest that grid-search-tuned algorithms can recover the main sleep–wake structure across extended recordings, even when the reference is a consumer wearable rather than PSG. However, smartwatch-derived labels should be interpreted cautiously, as consumer devices remain less reliable than PSG and vary across devices and populations (Lee et al., 2025).

Accordingly, the smartwatch analysis should be interpreted not as clinical validation but as a real-world stress test of longitudinal behavior. The goal is to evaluate whether tuned algorithms produce coherent multi-day sleep structure and plausible fragmentation patterns under realistic monitoring conditions. From this perspective, the single-subject analysis demonstrates the framework’s applicability in settings where wearable data are abundant but PSG is unavailable.

Fragmentation analysis further illustrates the limitations of movement-based sleep scoring. Although the algorithms consistently detected the main sleep episode, they differed in sensitivity to brief awakenings. Such awakenings often produce minimal motion, making them difficult to detect reliably. Algorithms with stronger smoothing may suppress these events, whereas more reactive algorithms may overcall wake. Parameter optimization alone cannot fully resolve this trade-off because fragmentation detection reflects the inherent balance between responsiveness and stability within each algorithm. This aligns with evidence that actigraphy-derived measures of wake after sleep onset and quiet wake are generally less reliable than total sleep time or sleep period time (Conley et al., 2019).

Several limitations should be acknowledged. First, the framework optimizes agreement among existing algorithms, so performance is constrained by their shared biases. Second, physiological plausibility constraints such as sleep-percentage bounds encode prior assumptions that may vary across populations or study designs. Third, the manual baseline reflects realistic exploratory practice but did not involve multiple raters, so comparisons between manual and automated tuning should be interpreted as proof of principle. Finally, the multi-day dataset involved a single participant with a smartwatch-derived reference, limiting generalizability.

Future work should evaluate the framework across larger and more diverse cohorts, including populations with insomnia, fragmented sleep, circadian disruption, or neurological disorders. Additional unsupervised objectives—such as transition consistency, day-to-day regularity, or alignment with behavioral markers like sleep diaries, light exposure, or heart-rate-derived sleep periods—could also be incorporated. Extending the ensemble approach toward weighted or probabilistic consensus may also improve robustness.

Overall, this study shows that consensus-based grid search provides a practical and scientifically defensible approach for tuning classical actigraphy algorithms when PSG is unavailable. Although it cannot overcome the intrinsic limitations of accelerometry for detecting quiet wake or fragmentation, it offers clear advantages over informal manual tuning in transparency, reproducibility, scalability, and consistency. The ensemble outputs further enhance robustness by summarizing shared structure across algorithms rather than relying on a single method. In this sense, the framework functions not as a new sleep detector but as a calibration and harmonization layer that improves the usability of existing algorithms in real-world, label-sparse actigraphy research (Patterson et al., 2023).

## 6. Declaration of AI and AI-Assisted Technologies

The conceptualization, core logic, and initial codebase of this project were developed solely by the author. AI-assisted tools (specifically GitHub Copilot, Claude, and Gemini) were employed for the following purposes:

### Code Optimization

Refining scripts and verifying algorithmic integrity against mathematical formulas and established open-source packages.

### Language Refinement

Improving grammatical structure, clarity, and technical writing style.

### Critical Review

Simulating a peer-review perspective to identify potential weaknesses in the manuscript’s logic and presentation.

In all instances, these tools were utilized under strict privacy settings to ensure that no part of the conversation or data was used for further model training. The final intellectual content and conclusions remain the sole responsibility of the author.

## 8. Code availability

GitHub: git@github.com:arahjou/GridSearch_Actigraphy_Optimization.git

---

**Supp 1.**
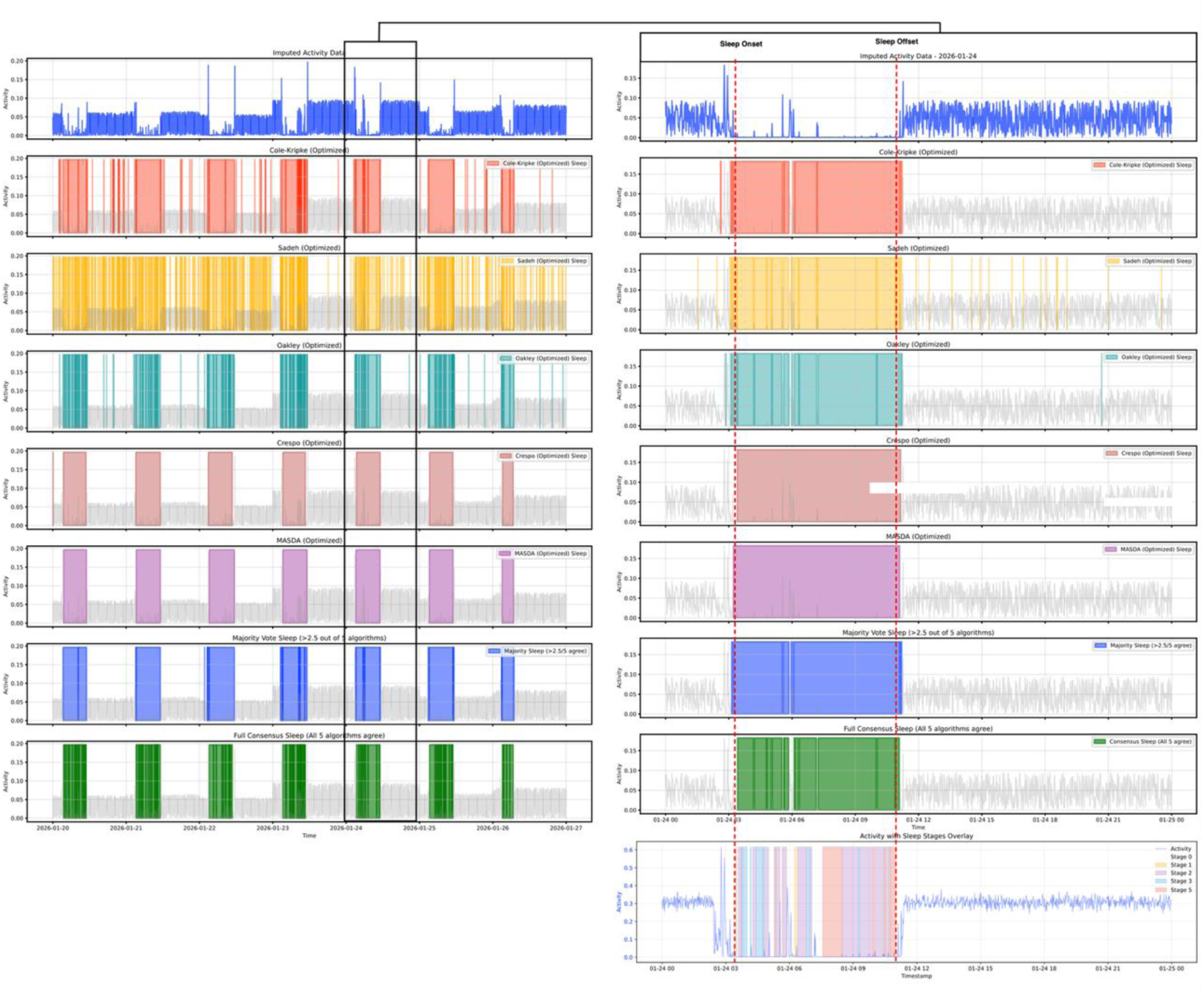
Sleep algorithm performance against Polysomnography (PSG) ground truth. (Left) Activity profile and algorithmic sleep classifications for a single subject over a 7-day period from the PSG validation dataset. The top row displays the raw mean absolute deviation (MAD) activity signal. Subsequent rows show binary sleep predictions (colored blocks) for four optimized algorithms (Cole-Kripke, Oakley, Crespo, MASDA/MASDA) and two ensemble methods. Note that the Sadeh algorithm was excluded from this analysis due to convergence failures. (Right) Detailed 24-hour view of a single recording day. Vertical red dashed lines indicate Sleep Onset and Sleep Offset. The bottom-most row displays the concurrent PSG hypnogram (Sleep Stages: N1, N2, N3, REM), serving as the reference standard. The visualization highlights the “quiescence vs. sleep” discrepancy discussed in the text: while all algorithms and ensembles successfully identify the broad sleep window (high inter-algorithm agreement), they produce solid blocks of sleep prediction that often fail to capture the physiological wake transitions and sleep architecture nuances present in the PSG recording.

**Supp 2.**
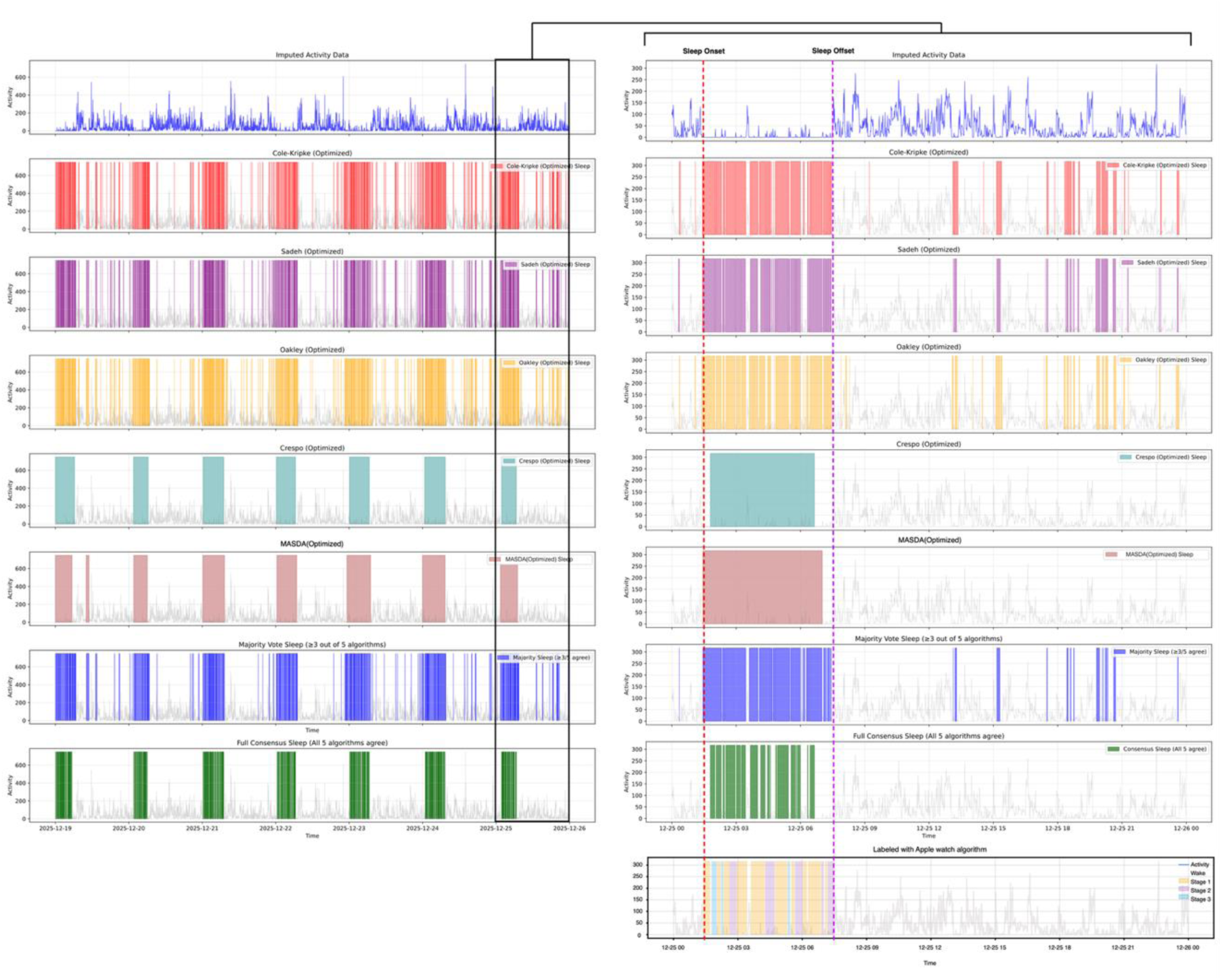
Multi-algorithm sleep detection and ensemble convergence compared to Apple Watch reference. (Left) Full 10-day activity profile (top row, blue) and corresponding binary sleep classifications generated by five optimized algorithms (Cole-Kripke, Sadeh, Oakley, Crespo, MASDA) and two ensemble strategies (Majority Vote, Full Consensus). (Right) Detailed 24-hour view of a single night (highlighted by the black bounding box) visualizing algorithmic behavior during sleep transitions. Vertical dashed lines indicate Sleep Onset (red) and Sleep Offset (purple) derived from the Apple Watch reference. The comparison highlights the structural differences between algorithms: threshold-based methods (Cole-Kripke, Sadeh, Oakley) exhibit high fragmentation and sensitivity to brief movements, while smoothed approaches (Crespo, MASDA) produce consolidated sleep blocks but may miss short awakenings. The bottom row displays the concurrent Apple Watch sleep staging (Wake, Light, Deep, REM) used as the validation standard for this dataset.

**Supp 3.**
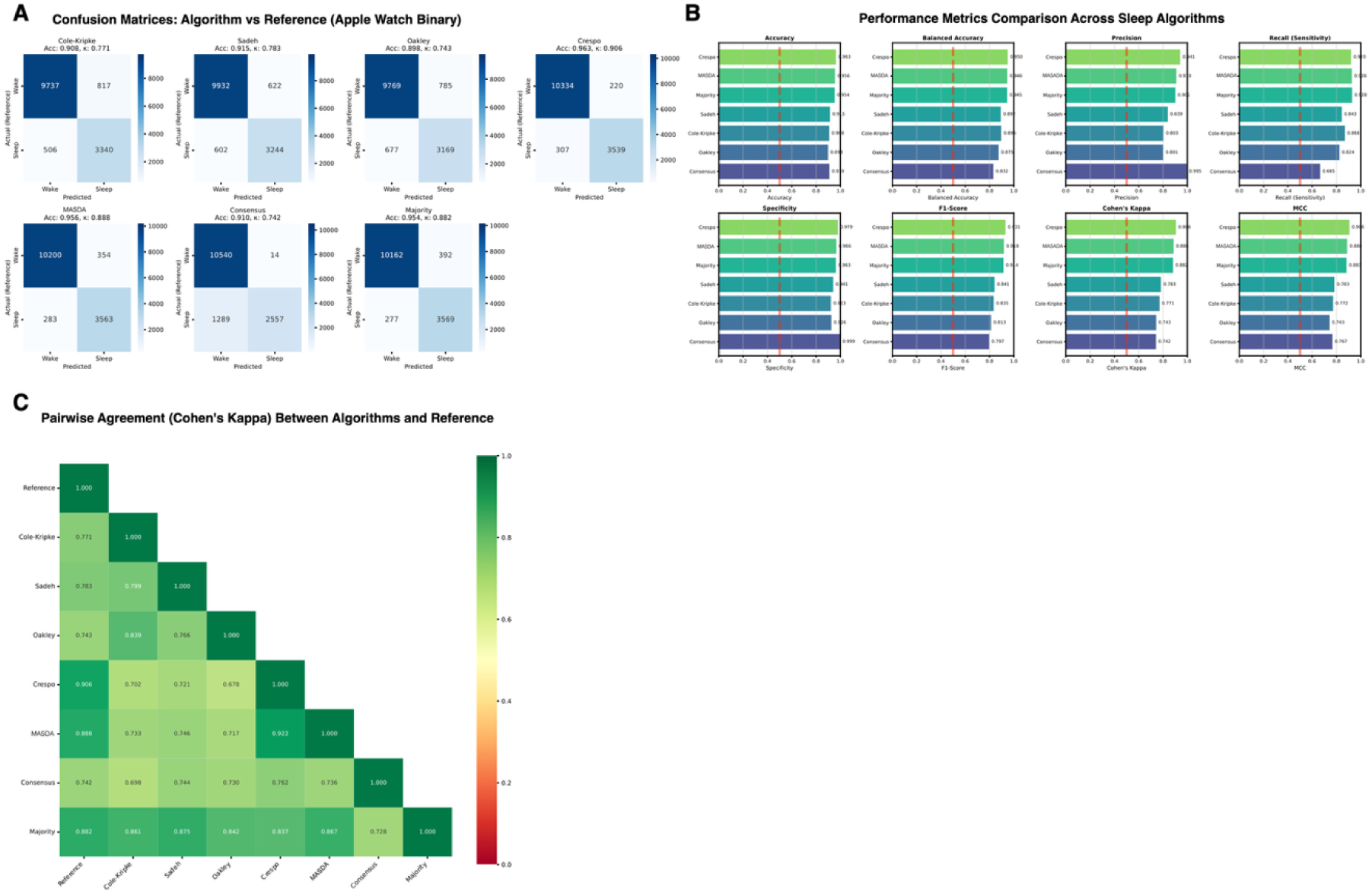
Multi-day sleep–wake detection performances The subject wore a research actigraphy device and an Apple Watch Series 10 simultaneously across multiple consecutive days. Apple Watch sleep staging was converted into binary sleep–wake labels (sleep = 1, wake = 0) and used as the reference for evaluation. (A) Continuous multi-day activity recording showing actigraphy signal and corresponding sleep–wake classifications produced by the evaluated algorithms using grid-search optimized parameters. The Apple Watch–derived binary sleep–wake labels are shown as the reference. (B) Multi-day comparison between algorithm predictions and the reference sleep–wake labels. Periods of sleep fragmentation are highlighted where repeated wake transitions occur within the sleep interval. (C) Quantitative evaluation of algorithm performance across multiple nights, including standard classification metrics (e.g., accuracy, sensitivity, specificity) computed relative to the Apple Watch reference.

